# A spiking neural network model of the Superior Colliculus that is robust to changes in the spatial-temporal input

**DOI:** 10.1101/2021.10.05.463158

**Authors:** Arezoo Alizadeh, A. John Van Opstal

**Affiliations:** Dept. Biophysics, Donders Centre for Neuroscience, Radboud University Heyendaalseweg 135, 6525 EZ Nijmegen, The Netherlands

**Keywords:** Saccades, Motor map, Midbrain, Neural encoding, Neural population, Primate

## Abstract

Previous studies have indicated that the location of a large neural population in the Superior Colliculus (SC) motor map specifies the amplitude and direction of the saccadic eye-movement vector, while the saccade trajectory and velocity profile are encoded by the population firing rates. We recently proposed a simple spiking neural network model of the SC motor map, based on linear summation of individual spike effects of each recruited neuron, which accounts for many of the observed properties of SC cells in relation to the ensuing eye movement. However, in the model, the cortical input was kept invariant across different saccades. Electrical microstimulation and reversible lesion studies have demonstrated that the saccade properties are quite robust against large changes in supra-threshold SC activation, but that saccade amplitude and peak eye-velocity systematically decrease at low input strengths. These features are not accounted for by the linear spike-vector summation model. Here we show that the model’s input projection strengths and intra-collicular lateral connections can be tuned to generate saccades that follow the experimental results.

**Author statement:** The midbrain SC generates fast saccadic eye movements through a large population of cells within a topographically organized motor map, in which the location, spike count and temporal firing patterns of recruited cells determine saccade metrics and kinematics. According to the dynamic ensemble-coding model, each recruited SC cell contributes to the saccade by linear vector summation of all its spike contributions. Our previous spiking neural network model used invariant cortical inputs to the SC cells for all saccades. We here improved the robustness of the model to large spatial-temporal variations in the input patterns, by tuning its top-down and lateral synaptic connections, to generate saccades with properties observed in electrophysiological experiments.

## 1. Introduction

### Background

Saccades are fast eye movements that redirect the fovea to a peripheral target. They obey a stereotyped ‘main-sequence’ kinematic relationship between saccade amplitude and movement duration (an affine relation) and between amplitude and peak eye-velocity (a saturating function; [1]). As the duration of the acceleration phase is roughly constant across a wide range of amplitudes, saccade-velocity profiles are positively skewed, whereby skewness increases with saccade duration [2]. Moreover, because trajectories are nearly straight, the horizontal and vertical saccade-velocity profiles are scaled versions of each other, thereby approximately matched in duration and shape [3-6].

Together, these kinematic features betray nonlinear processing in the generation of saccades. In earlier models of saccade control, the saturation of peak eye velocity was believed to reside in the (passive) saturation of firing rates of brainstem pre-motor burst neurons [7-8]. Later studies, however, have suggested that these properties may instead betray a deliberate optimal control strategy that aims to optimize speed-accuracy trade-off in the presence of multiplicative and additive neural noise [9-13]. Single-unit recordings and quantitative modelling of instantaneous spiking behavior of saccade-related cells in the midbrain Superior Colliculus (SC) have suggested that such a mechanism might be implemented at this oculomotor midbrain level [13-15].

The SC is a primary source of gaze-motor commands to the brainstem saccade generators [8, 14, 16-21], and is recruited for all voluntary and involuntary saccades. Its deeper layers contain an eye-centered topographic map of visuomotor space [16, 19, 22], in which the location and total spike count of the neural population encode the saccade amplitude and direction [17-19]. Several studies have suggested that the temporal firing profiles of the neural population may also specify the instantaneous saccade trajectory and its velocity profile [13-14, 23-26].

Although also the frontal eye fields (FEF) and posterior parietal cortex (PPC) are strongly involved in saccades, their major role appears to be in the preparation of higher-level, reward-contingent, and task-relevant eye-movements, like anti-saccades [27], target selection and identification [28], saccade suppression [29], also when its planning is already in progress [30], or towards remembered targets [31-32]. Their main outputs are transferred to the SC, which thus constitutes a final common pathway for saccade initiation and control.

Schiller and colleagues examined the effects of FEF and SC ablations on eye movements [33]. The deficits caused by a lesion of either structure appeared to be rather subtle when monkeys were tested a few days later and recovered over time. However, when both structures were removed, monkeys were no longer able to redirect their gaze to peripheral targets. In contrast, Hepp et al. reported a strong reduction (near-abolition) in frequency and velocity of visual-evoked spontaneous saccades and quick phases of vestibular nystagmus immediately following bilateral muscimol-induced SC inactivation, indicating a crucial role for the SC output to voluntary and involuntary saccades [34]. Also, acute FEF inactivation strongly affects the properties of visual-evoked saccades [31-32]. Thus, the immediate effects of SC and FEF inactivation seem to be much stronger than seen with the earlier longer-term ablation studies [28, 33]. Presumably, the FEF can take over SC function during the recovery period, when the latter is no longer available.

Recently, Peel and colleagues examined the acute influence of inactivating FEF by local cooling on saccade metrics and kinematics, and on the associated neural firing patterns of saccade-related SC cells for different saccade tasks [35]. Their results indicated that FEF inactivation did not significantly affect direct visual-evoked saccades but led to a significant decrease of about 10% in SC spiking activity for memory-guided saccades. The authors suggested that these cortically mediated saccades may utilize, besides the direct FEF-SC-brainstem pathway, an additional, flexible processing circuit that bypasses the SC.

### Problem statement

In the present paper we focused on the encoding of saccades, generated by the direct cortical-SC-brainstem pathway. Single-unit recordings of saccade-related cells in the SC have indicated that the peak firing rate, burst duration, and shape of the burst profile of the central neuron in the population depend systematically on its location in the map [14]. Moreover, each SC neuron elicits about a fixed number of spikes for its preferred saccade, irrespective of its motor map location.

These features were incorporated in a simple neuro-computational feedforward spiking neural network model, in which each spike of each recruited neuron encodes a fixed (tiny) movement contribution to the saccade that is solely determined by its location (the cell’s ‘spike vector’). The saccade trajectory then results from dynamic linear summation of all spike vectors from the spike trains from all cells in the population. Because linear spike-vector summation, in combination with a linear brainstem model could reproduce the full repertoire of (nonlinear) saccade kinematics and their trajectories, we argued that the firing patterns within the SC motor map were responsible for the nonlinear main-sequence properties, velocity profiles, and component cross-coupling of saccades [15]. The SC motor map would thus embed an optimal control for saccade generation.

Electrical microstimulation in the SC has revealed that the evoked saccade amplitude varies systematically with the applied current strength: at low currents, amplitudes are small, increasing to a site-specific maximum at higher current strengths, determined by the electrode’s position in the motor map [36-38]. In addition, small saccades evoked at low intensities are also slower than visual-evoked main-sequence saccades of the same amplitude. Further, variation of the stimulation pulse rate systematically affects the eye velocity ([37], in monkey; [39], in barn owl). So far, these input-dependent properties had not been accounted for by the linear ensemble-coding model [40].

### This study

We here extend the spiking neural-network model of [40-41] with the aim to increase its robustness to large variations in spiking input patterns arising from cortical sources. To simplify the analysis, we constructed a one-dimensional model with a cortical input layer and a collicular output layer. We independently varied the input spiking patterns in the spatial (i.e., population extent) and temporal (burst durations and peak firing rate) domain, reminiscent to the presumed effects of electrical stimulation, partial inactivation, or visual stimulation at different intensities, and input stimulus durations. We re-tuned the intra-collicular excitatory-inhibitory synapses and top-down connections, with the aim to generate saccades with similar input-dependent metrics and kinematics as has been observed in electrophysiological experiments.

## 2. Methods

### 2.1 Network architecture

We constructed a two-layer spiking neural network model with a cortical input layer, and a layer of SC output neurons, respectively (Fig 1). Each layer consists of 200 neurons, uniformly distributed on 0-5 mm of the horizontal meridian of the SC motor map. In the linear dynamic ensemble-coding model [13-14], the saccade kinematics are fully determined by dynamic cumulative summation of all spike vectors in the neural population during the saccade. The saccade trajectory is thus generated by:

**Fig 1.**
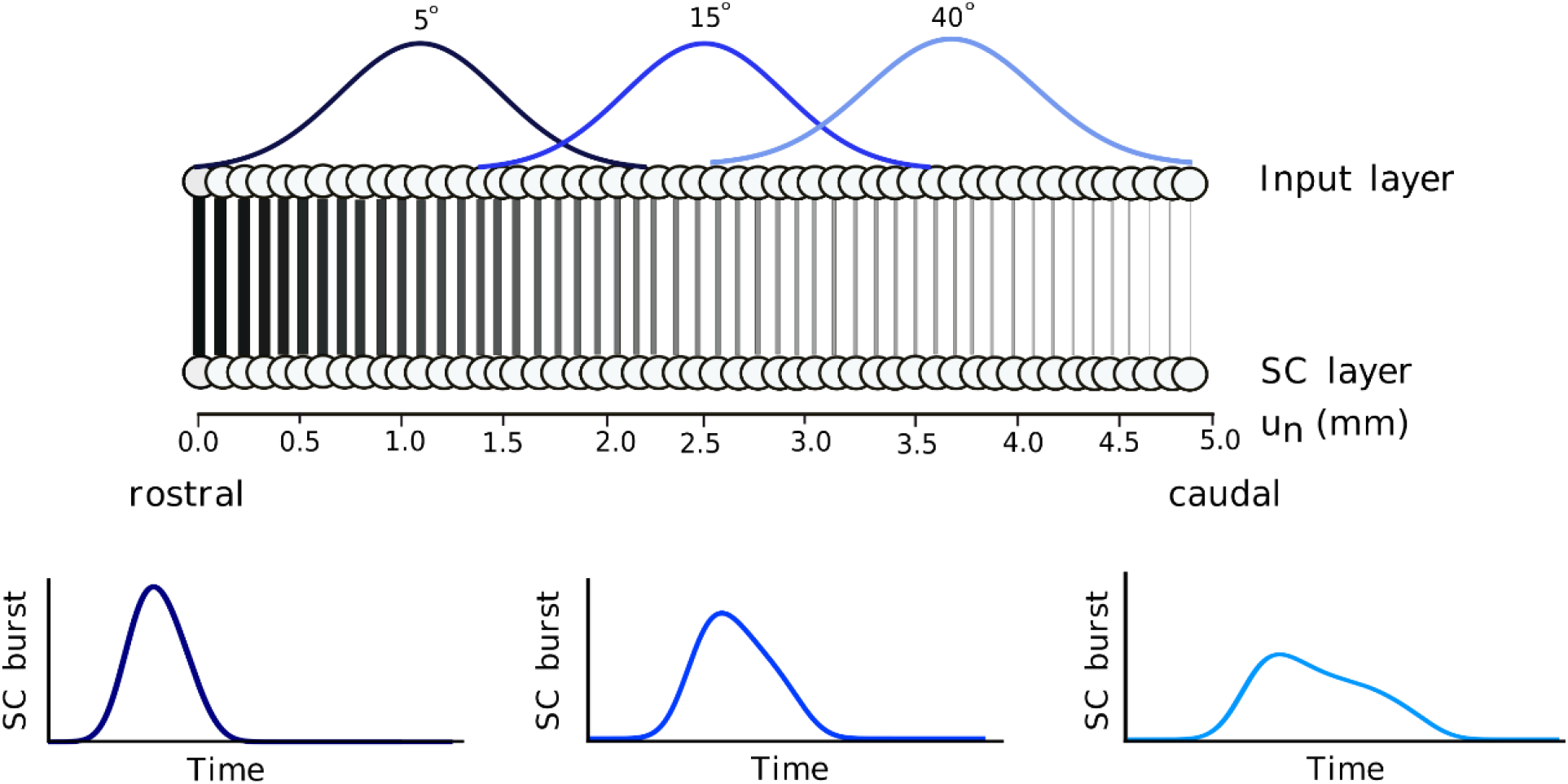
Schematic overview of the two-layer feedforward neural network. The spiking neural network model generates different saccade-related bursts (bottom) that are evoked by a spatially translation-invariant input population (top), here positioned at T=5, 15 and 40 deg eccentricity. Thickness of the lines for the downward projections symbolize the synaptic connection strengths, 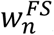, between the input and SC layers (high for the rostral zone, low for the caudal zone; Eqn. 6).

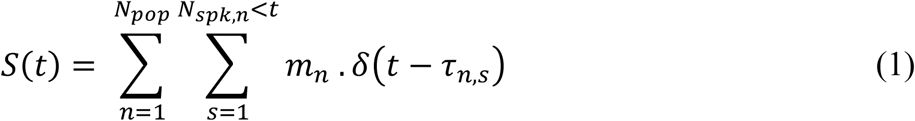

in which *δ*(*t* − *τ*_*n,s*_) is a spike of neuron *n* at time *τ*_*n,s*_, *N*_*pop*_ is the total number of active neurons in the population, *N*_*spk,n*_ is the total number of spikes fired by neuron *n*, and *m*_*n*_ is the cell’s site-specific, fixed ‘spike vector’. The latter is determined by each neuron’s efferent synaptic connection strength to the horizontal brainstem circuitry [19], which we here simply specified in one dimension as:

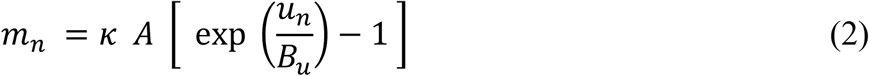

with *u*_*n*_ ∈ [0 − 5] mm the cell’s anatomical rostral-caudal location in the SC map, encoding the horizontal saccade amplitude, and *κ* a fixed scaling factor that only depends on the assumed cell density (in number neurons/mm). This scaling factor was calibrated for a horizontal saccade of 15 deg. The SC afferent mapping parameters (here: *A*=3.0 deg, and *B*_*u*_=1.4 mm) had been estimated from [16, 19].

The input layer receives an external input signal from other (unspecified) inputs, which it transforms into spiking activity through its neural dynamics (all neurons in the model are governed by the AdEx neural model equations; see Supporting Information; [41]). For simplicity, the input-layer neurons do not interact with each other. The input-layer spikes are subsequently transmitted by topography-preserving one-to-one synaptic connections to the neurons in the SC layer. The biophysical parameters of the SC neurons, such as their adaptation time constant, their synaptic connection strengths with the input layer, and their lateral excitatory-inhibitory connections, depend on their location in the motor map, and as a result identical firing rates in the input layer at different locations will lead to dissimilar responses of the SC cells (Fig 1, bottom). As described below, the network is tuned such that these responses, and the ensuing saccade (Eq. 1) follow similar characteristics as observed in the electrophysiological recordings and microstimulation experiments.

We simulated eye movements in response to the presentation of a single input stimulus at a particular location in the input layer. A target point in visual space at *T* deg from the fovea was thus mapped to its anatomical position, *u*_*T*_, corresponding to the center of a Gaussian-shaped population in the input layer according to the afferent mapping function of Ottes et al. ([19]; Fig 1, top):

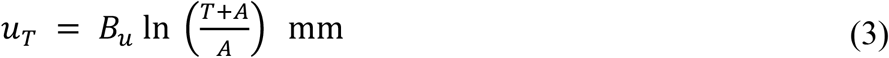

The one-dimensional model was simulated with the Brian2 spiking neural network simulator [42]. We modeled the neurons in the network by the adaptive exponential integrate-and-fire (AdEx) neuron model [43], as the parameters of this model can be readily related to physiological quantities. Details of this neural model, including the chosen parameter values, are provided in the Supporting Information, S1 Table, and [41]. Here, we only highlight the major differences with the earlier model.

### 2.2 External input current

We provided an external input current to the network around the image point, *u*_*T*_, of the desired target, *T*, in the input layer, leading to an input population spiking activity centered around the image point, *u*_*T*_ (Eq. 3; Fig 1, top). The central neuron in the input population receives the maximum input activation current, *I*_*0*_*(t)*, while the other neurons in the input layer are stimulated by current strengths that decay as a Gaussian with distance from *u*_*T*_. The spatial-temporal external input current was thus described by a separable spatial-temporal function on the input neurons by:

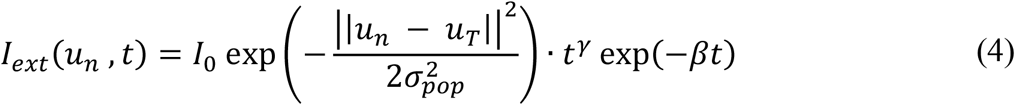

where *u*_*n*_ is the anatomical position of a neuron in the input map, *σ*_*pop*_ determines the size of the input population, *t* is time (s), *u*_*n*_ is the location of neuron *n* (mm), and *I*_*0*_ is the maximum input amplitude (pA). The time-dependent term is a gamma function, characterized by γ (skewness, dimensionless) and β (measure for inverse duration, in s^-1^).

To investigate the relationship between the resulting saccade metrics, trajectories, and kinematics as function of the input current profiles, we varied the input current in both the spatial and the temporal domain. The default input stimulation profile (serving as the model’s control condition) was defined by the following parameters: I_0_ = 3.0 pA, σ_pop_ = 0.5 mm, β = 0.03 s^-1^, and *γ*=1.8.

#### Spatial input variation

In the spatial simulations, we varied the stimulated input population size between σ_pop_ = 0.05-1.0 mm. Input amplitude varied between I_0_ = 2.0-3.0 pA for input population size lower than 0.5mm and it is constant at 3.0 pA for input population size higher than 0.5mm. The temporal stimulation parameters were kept fixed at their default values: β = 0.03 s^-1^ and *γ*=1.8. This parameter variation led to the activation of 10 - 200 input layer neurons (e.g., Fig 2A, B).

**Fig 2.**
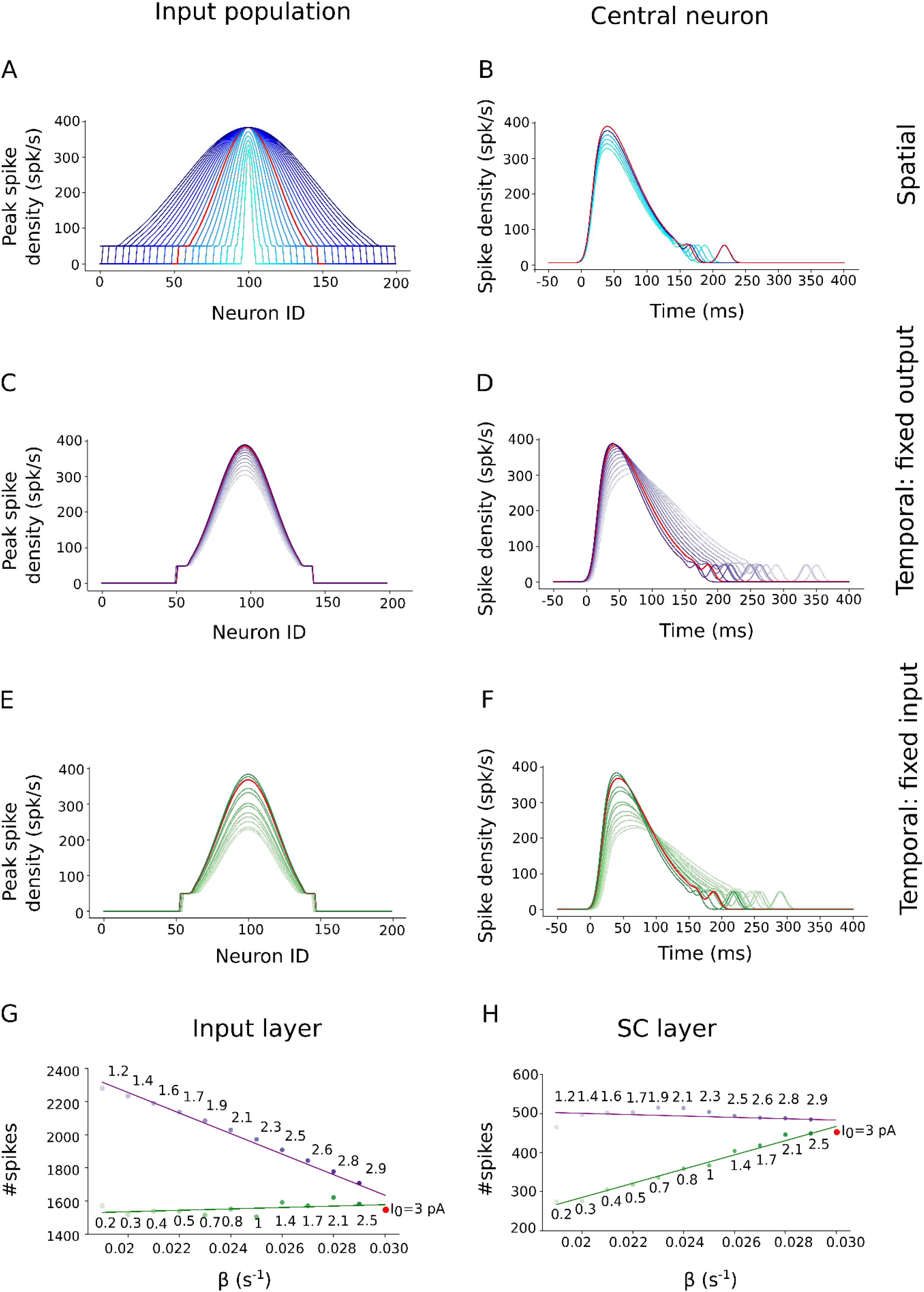
(A-F) Burst profiles of the input layer neurons in response to changes in the spatial-temporal parameters of the external input currents (see also S1 Table in Supporting Information). Red curve corresponds to the default parameter set. (A) Spatial distribution of the peak spike-density functions for σ_pop_ in 0.05-1.0 mm, and I_0_ in 2.0-3.0 pA at T=15 deg. (B) Spike densities as function of time for the central neuron of the populations in (A). (C, D) Spatial (C) and temporal (D) spike-density distributions for β in 0.019-0.03 s^-1^, and I_0_ in 1.2-3.0 pA at the T=15 deg site (σ_pop_=0.5 mm). Parameter values of the input currents were chosen such that the SC output generated a fixed number of spikes. (E, F) Spatial (E) and temporal (F) spike-density distributions for β in 0.019-0.03 s^-1^ and I_0_ in 0.2-3.0 pA. Parameters now ensured a fixed number of spikes in the input layer. (G) Total number of spikes in the input layer for the two temporal variation scenarios (green and purple data). (H) Same as in (G) for the output layer.

#### Temporal input variation

To investigate the influence of input spike rates (firing frequency) to the SC motor output and SC population activity, we also varied the input current in the temporal domain. In these simulations, the externally applied input current always activated a fixed population size of σ_pop_ = 0.5 mm. We then considered two different scenarios:

i. A variable stimulation duration (β) between 0.019-0.030 s^-1^, and a current intensity (I_0_) between 0.2-3.0 pA, but selected such that the resulting number of spikes in the SC *output layer* remained fixed (e.g., Fig 2C, D and Fig 2G, H (purple)).
ii. A similar variation in the input, but now such that the number of spikes sent from the *input layer* to the SC remained constant: β varied between 0.019-0.03 s^-1^, and I_0_ between 1.2-3.0 pA (e.g., Fig 2E, F and Fig 2G, H (green)).

Fig 2 illustrates these different external stimulation input scenarios. The first three rows of the left-hand column (Fig 2A,C,E) show the spatial distributions of the peak firing rates of the neurons in the input population, when the target stimulation point corresponded with T = 15 deg (i.e. at *u*_*T*_ = 2.5 mm); the right-hand column (Fig 2B,D,F) shows the temporal profiles of the spiking patterns for the central neuron in the input population at *u*_*T*_ = 2.5 mm. Fig 2A,B shows the spiking responses of the input layer neurons when the external input width was systematically varied between 0.05 and 1.0 mm, and the amplitude between 2.0 and 3.0 pA. The red curve corresponds to the default control stimulation (σ_pop_ = 0.5 mm). To avoid non-physiologically high firing rates, we imposed an upper limit to the evoked firing rates in the input layer (by including a saturating sigmoid input-output relationship) at 400 spikes/s.

Fig 2C, D shows the spike-density functions of the input layer when the input current at *u*_*T*_ = 2.5 mm stimulated a fixed population size (0.5 mm), but with a variable current duration (β) and strength (*I*_*0*_). These latter two parameters were selected such that the SC neural population generated a fixed number of output spikes (Fig 2H, purple line). Note that the amplitude of the spike density function of the central input neuron decreased with decreasing external current strength; at the same time, burst duration in(de-)creased with de(in-)creasing stimulus strength.

Fig 2E, F shows the input layer responses to the external currents when the input duration and strength were tuned such the input population sent a fixed number of spikes to the SC. Fig 2G, H illustrates how the number of spikes of the input- (G) and output (H) layers varies in response to the chosen input currents with variable temporal behavior: a constant number of spikes of the input (green), vs. output layer (purple). Note that if the number of spikes is constant in one layer, it either decreases (input) or increases (output) in the other layer.

### 2.3 Superior Colliculus cells

The neurons in the SC layer receive the total synaptic input current, given by the synaptical weighted sums of the spikes from the input-layer, and from the SC neurons themselves, whereby the latter are relayed by conductance-based lateral excitatory-inhibitory synapses (S4 Eq., Supporting Information). Because of the location-dependence of the parameters specifying the AdEx equations for the SC neurons, their activity patterns depend on their location in the motor map. Neurons near the rostral site generate a small saccade with a high-frequency, short-lasting burst of activity, while at caudal sites the evoked activity has a lower peak firing rate, and longer burst duration, associated with a large saccade.

#### Lateral intra-collicular connections

The saccade-related neurons in the SC population communicate with each other through lateral interactions, which cause all bursts to approximately synchronize with the central cell. In the original version of the model, these interactions were described by a “Mexican-hat” function (short-range excitation, and long-range inhibition [44]), which acts as a soft winner-take-all mechanism [41].

Two Gaussians describe the spatial extent of the excitatory and inhibitory connection profiles, between a neuron, *n*, and any other neuron, *i*, in the motor map (apart from itself) as function of anatomical position. In the present study, we slightly modified the earlier proposal to:

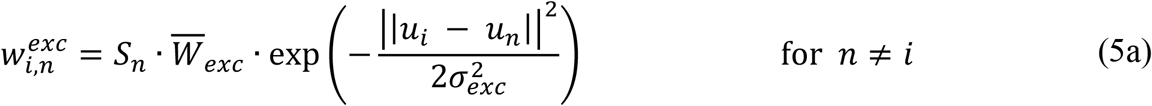

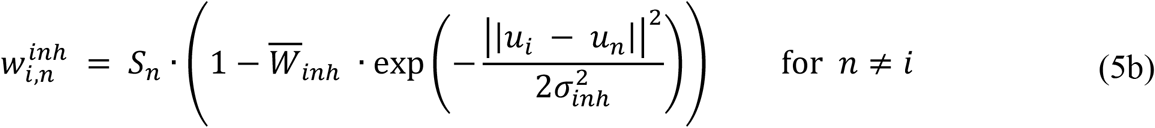

with 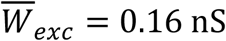 and 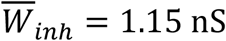 fixed excitatory and inhibitory weight parameters. The location-dependent gain, *S*_*n*_, causes the lateral interaction scheme to be site-dependent. These lateral connections have a direct effect on the spiking behavior of each neuron, and hence on the overall network dynamics. Strong excitation (re. inhibition) would result in an unbounded spread of the population activity across the motor map (and hence, an ever-increasing saccade amplitude), whereas strong inhibition would quickly fade out the neural activity altogether. We aimed to find parameter values that would ensure a balanced amount of excitation and inhibition, leading to a stable Gaussian population activity, in such a way that (considerable) spatial-temporal changes in the input population activity (as illustrated in Fig 2) would lead to experimentally observed changes in the SC output saccades (Eqn. 1).

### 2.4 Network tuning

We employed brute-force search algorithms to find suitable values for the lateral inhibitory and excitatory weight parameters, the feedforward projection strengths from input to output layer, and for intrinsic properties of the AdEx equations of the SC neurons.

Besides 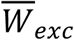 and 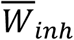, we also tuned the widths of the Mexican-hat profiles (σ_inh_ and σ_exc_) to yield an appropriate SC population size with synchronized activity, also when the total input activity profile would far exceed the normal default size of 0.5 mm (Fig 2A). We further extended the model with the lateral synaptic gain parameter, *S*_*n*_, as a location-dependent excitatory and inhibitory scaling.

The intrinsic biophysical parameters of the AdEx equations for the SC neurons (Supporting Information) were optimized by systematically varying their adaptation time constant, τ_q,n_, in combination with the location-dependent feedforward synaptic projection strengths between the layers, 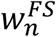. In addition, we assessed the effects of varying the location dependence of the intra-collicular scaling parameter, *S*_*n*_, on the saccade trajectories.

The adaptive time constant of the neurons affects the susceptibility of the neuron to synaptic input and influences their instantaneous firing rate and bursting properties, and hence the kinematics of the saccade. As the feedforward synaptic projection strength between the input layer and SC layer determines the number of presynaptic spikes that is transferred from the input layer to the different locations of the SC layer, it mainly affects the SC neuron’s peak firing rate. The intra-collicular synaptic gain, *S*_*n*_, normalizes the SC output against variability in the total input activity. Together, these three parameters caused a systematic change in the firing properties of SC cells along the rostral-caudal axis of the motor map, while ensuring a fixed maximum number of spikes for the neurons’ preferred saccades, *N*_*u*_*(R)*, with a sigmoid-like response sensitivity to large changes of the input firing patterns.

We employed a similar brute-force search [40-41] to find the optimal location-dependent values of 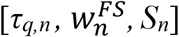 that ensured a fixed number of spikes per neuron for a saccade that kept a constant amplitude and peak velocity for input patterns far exceeding the default strength of *σ*_*pop*_ = 0.5 mm (e.g., Fig 2A). Note that the input currents in which we also varied the temporal stimulation properties (β) were not used to tune the parameters of the network.

Equation 6 explains the results of the network tuning for the adaptation time constant, *τ*_*q,n*_, and for the top down-projection strengths, 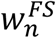, as function of the map coordinate, *u*_*n*_. Interestingly, to obtain appropriate saccade responses (see below), both parameters resulted to co-vary in a linear way with the anatomical rostral-caudal location:

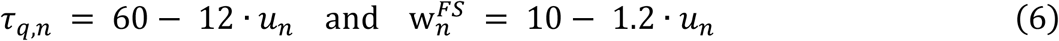

Fig 3A depicts the net intra-collicular lateral connection strengths from each neuron as obtained from the brute-force search. The lateral connections yield short-range excitatory and long-range asymmetric inhibitory effects from each neuron in the map. Effectively, SC neurons receive both excitatory and inhibitory potentials from cells endowed with different adaptation time constants, firing rates, and reversal potentials (Supporting Information, S1 Table). Due to the strong symmetric lateral inhibitory connections in the SC layer the number of active neurons in the SC layer saturates when the external input current causes a large recruited input population that may far exceed the standard size of 0.5 mm.

**Fig 3.**
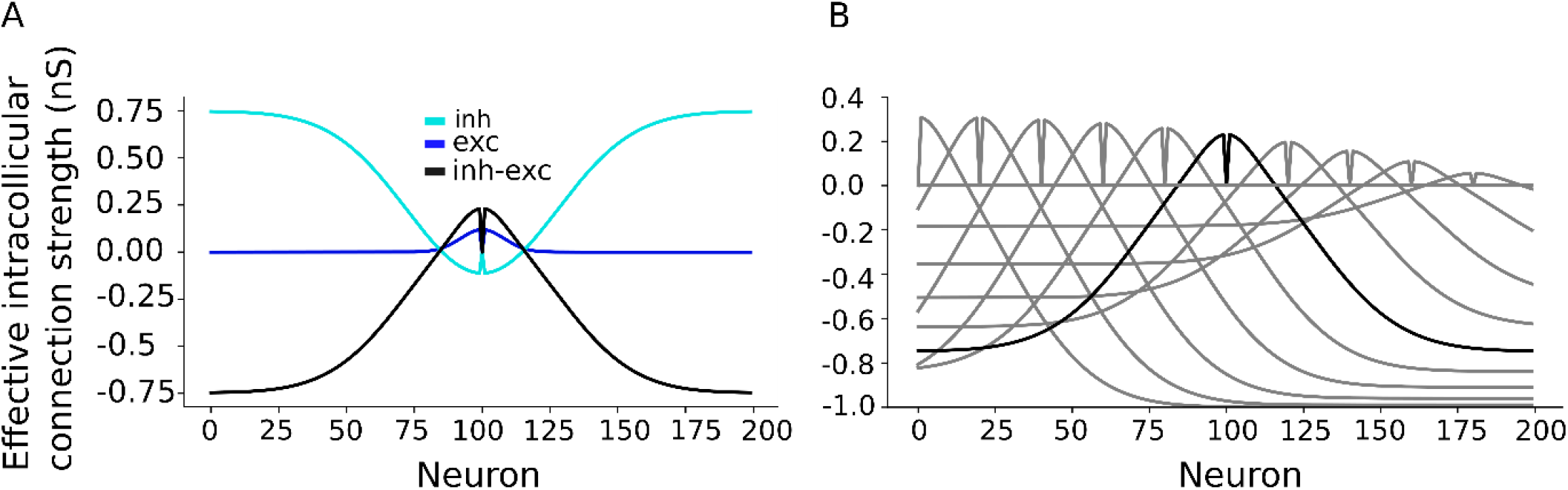
A) The excitatory (σ_exc_ = 0.2 mm; dark-blue) and inhibitory (σ_inh_ = 0.7 mm; light-blue, shown inverted) intra-collicular synaptic connections, and their total effect (black line) for central neuron at neural population generating 15 deg saccade, result in a symmetric local excitatory and global inhibitory connectivity. The net excitation around the neuron (at 0) approaches the value of +0.23. B) Total effect of excitatory and inhibitory intra-collicular synaptic connections for neurons across motor map generating different saccade amplitude. The intra-collicular synaptic connections are stronger towards the rostral zone (thus counter-acting the higher firing rates) than towards the caudal zone (where cells have lower firing rates).

Fig 3B shows the total intra-collicular lateral connection strengths for neurons across rostral to caudal site of the motor map. The lateral connections inhibitory and excitatory strength decreasing from rostral to caudal zones by scaling S_n_ to influence the shape of the nonlinear main-sequence relationship of the model’s saccades between their amplitude and peak eye velocity. The following heuristic relation provided satisfactory results (see Results):

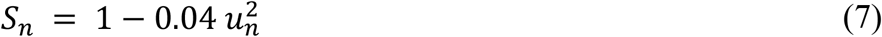

### 2.5 Eye-movement trajectories

Eye movements were encoded by the linear ensemble coding scheme of the population activity in the SC motor map (Eq. 1). We applied the one-dimensional efferent motor map of Eq. 2 to the new network configuration. The resulting eye-displacement vector, S(t), was smoothed with a Savitzky–Golay filter to compute the instantaneous eye velocity.

## 3. Results

### 3.1 Bursting behavior of SC AdEx neurons

To illustrate the effect of varying the input stimulation (Eqn. 4) on the response behavior of the AdEx model of a typical SC neuron, Fig 4 shows the time dependence of the neuron’s membrane potential, *V(t)*, when the input (applied at T=15 deg) varied in population size, σ_pop_ (Fig 4A), or in stimulus duration and intensity, β, *I*_*0*_ (Fig 4B).

**Fig 4.**
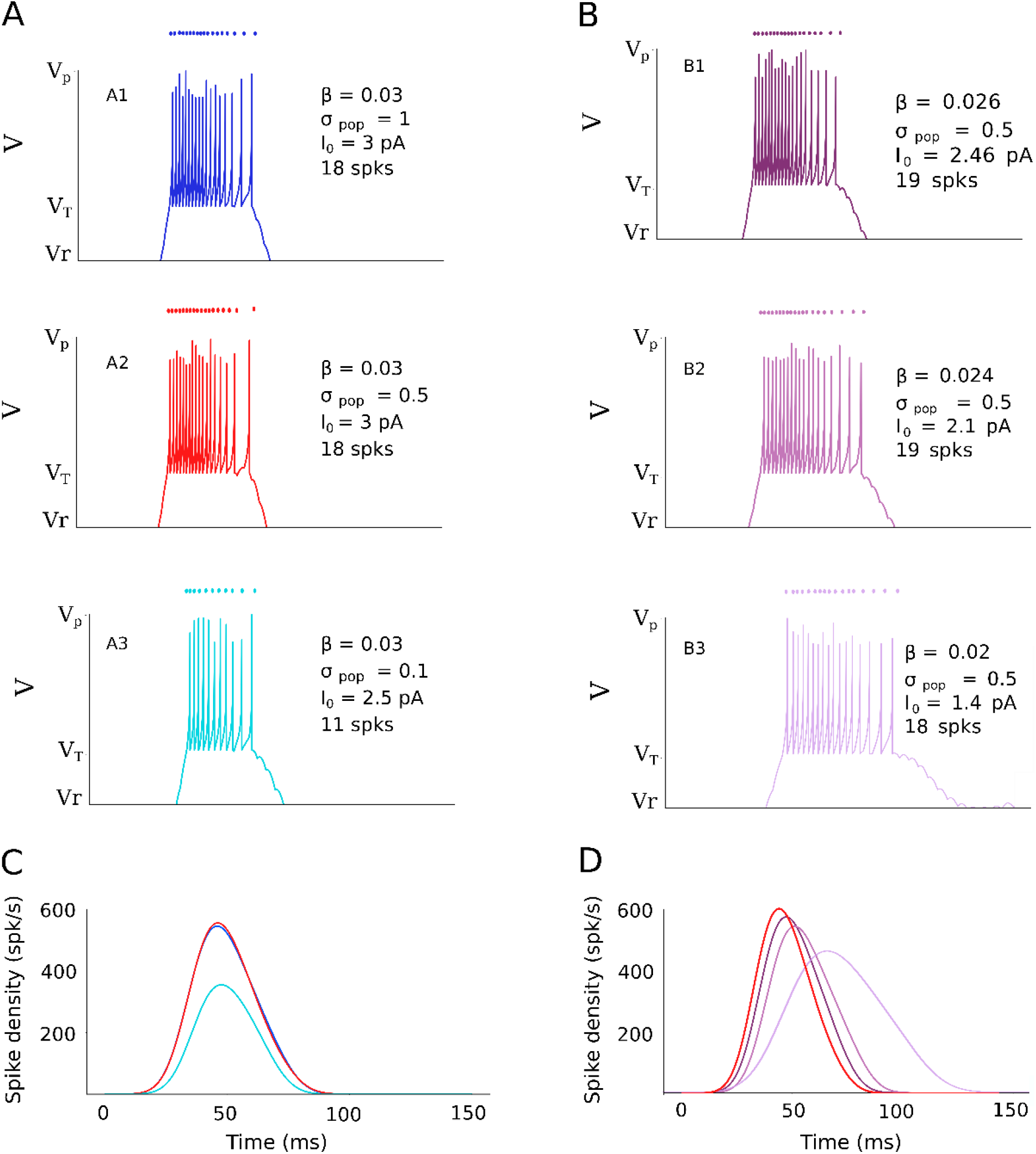
Effect of varying external current input parameters, such as the recruited input population size (σ_pop_), stimulus duration (β), and input stimulus amplitude (I_0_) on the bursting characteristics of an AdEx neuron at u_n_ = 2.5 mm in the SC output layer [τ_q_ =30, W ^FS^ =3]. Panels A,B show the membrane potential, V(t), and a dot-display of the individual spikes (top), for three input current population sizes around T=15 deg: (A1) σ_pop_ = 1.0 mm, (A2), σ_pop_ = 0.5 mm (the default stimulation input), (A3) σ_pop_ = 0.1 mm, and for three stimulus duration/intensity values: (B1) β = 0.026/I_0_=2.46, (B2) β = 0.024/I_0_=2.1, (B3) β = 0.020/I_0_=1.4. (C) Corresponding spike density functions for varying the external stimulation population size, and (D) for varying the external stimulus duration/intensity parameters. Line colors correspond to the traces in panels (A, B).

The SC neuron in the center of the population (at *u*_*T*_=2.5 mm) emitted fewer spikes, *N*_*spk*_ = 11, for the small input population size (σ_pop_ = 0.1 mm; A3), while it generated the same number of spikes, *N*_*spk*_ = 18, for the default size (σ_pop_ = 0.5 mm; A2) and for the much larger input population (σ_pop_ = 1.0 mm; A1). In all three cases, *V(t)* had the same duration, as the stimulation input current had the fixed default value of β = 0.03 s^-1^. The burst profiles of the neuron (Fig 4C) for the three different input currents had the same duration too, but the peak firing rate was clearly lower for the small input population size.

When the input current was given a fixed population size (0.5 mm) but varied in its duration and intensity parameters (Fig 4B), the resulting burst durations varied accordingly. The smaller β, the longer the membrane potential, *V(t)*, and, consequently, the resulting burst profile of the neuron. However, in all three cases the emitted number of spikes of the cell remained approximately constant at *N*_*spk*_ = 18 or 19.

### 3.2 Effect of lateral interactions

To establish the functional role of the intra-collicular lateral connections on network performance in response to large changes in the SC input patterns, we first look at the network’s behavior without these lateral interactions. Fig 5A illustrates how the number of active neurons in the SC layer depends on the input population size, when stimulation is applied at target location T=15 deg at different stimulation strengths, yielding different population widths (dark-green curve). The number of activated SC cells increases with the input population size up to a level where all 200 SC cells are recruited. Similarly, the total number of spikes emitted by the SC population appears to increase approximately linearly with the population size (light green). This means that the resulting saccade amplitude (Eq. 1), which was tuned to 15 deg for the default population size of 0.5 mm, keeps increasing to 27 deg for the input population size of 1.0 mm. Only the maximum number of spikes emitted by the central cell remains constant, because the input firing rates had been limited in the simulations to 400 spikes/s and central neuron receive input spike activity from only one neuron at input layer through one-to-one synaptic connections between two layers (see Methods, and Fig 2A).

**Fig 5.**
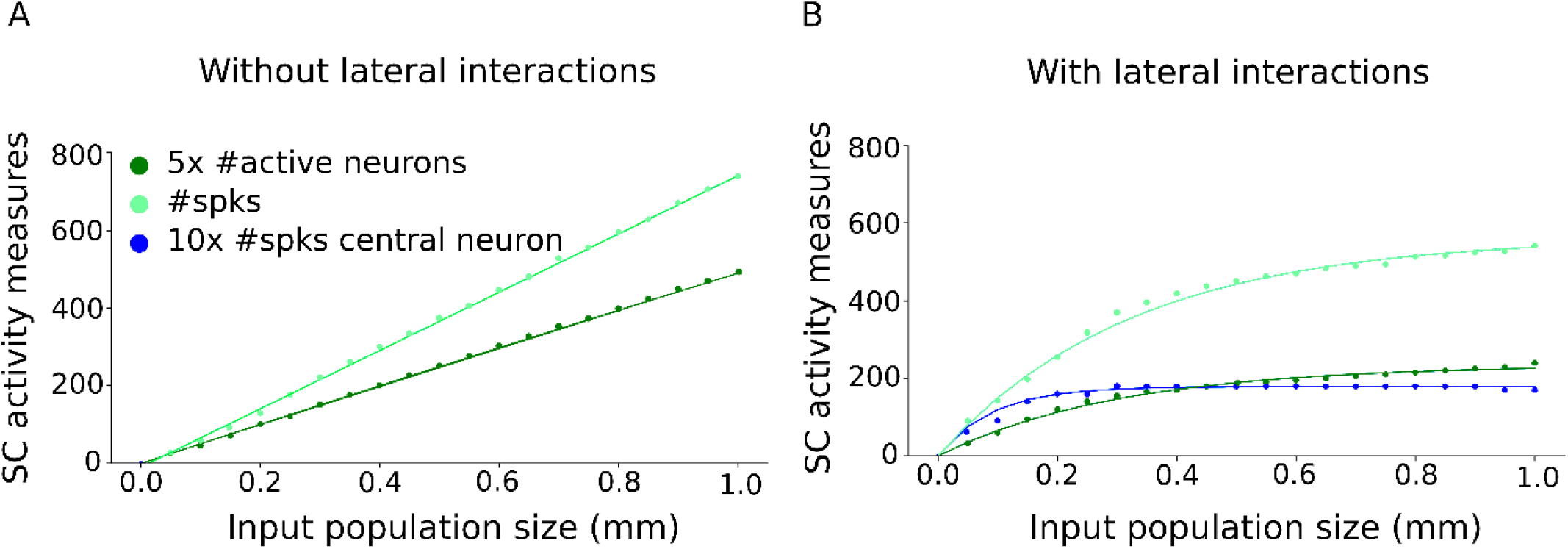
Number of active neurons (dark green) in the SC layer, the total number of spikes emitted by the neural population (light green), and the number of spikes emitted by the central cell in the population (dark blue) for a saccade towards T=15 deg, as function of the input population size: (A) in the absence of lateral connections, and (B) with the lateral connections specified by Eqns. 5-7. Without lateral connections, the total number of spikes and the number of activated SC neurons increase without bound (R=27 deg, V_pk_=1000 deg/s; not shown). Due to limiting stimulation strength to 400 spikes/s and considering one-to-one synaptic connections between two layers, number of central cell spikes does not increase by increasing external input current’s population size. With lateral interactions, the number of spikes from the population, and the recruited population size both saturate (R= 16.2 deg, V_peak_= 580 deg/s).

Fig 5B shows the results of the network model for the same input stimulation conditions, with the lateral interactions described by Eqns. 5-7. For small input populations (up to 0.3-05 mm), the number of recruited cells, the total number of spikes, as well as the number of spikes of the central neuron, all increase with the input population size. However, the number of activated neurons and the total number of spikes from the motor map now increases very little beyond the default stimulation strength at 0.5 mm. As a result, the saccade amplitude follows a saturating relationship with the input stimulation intensity (see also below).

### 3.3 Model output at default input

In Fig 6 we quantified the collicular bursts in response to the external input current with default parameters: *σ*_*pop*_ = 0.5 mm, *I*_*0*_ = 3.0 pA, and *β =* 0.03 s^-1^, applied at different sites in the input map. Appropriate tuning of the biophysical parameters of the SC cells, such as τ_q,n_ and 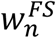 along the rostral-caudal axis in the SC map ensured burst profiles that faithfully reflect the experimentally observed spatial variations in the saccade-related SC bursts [13].

**Fig 6.**
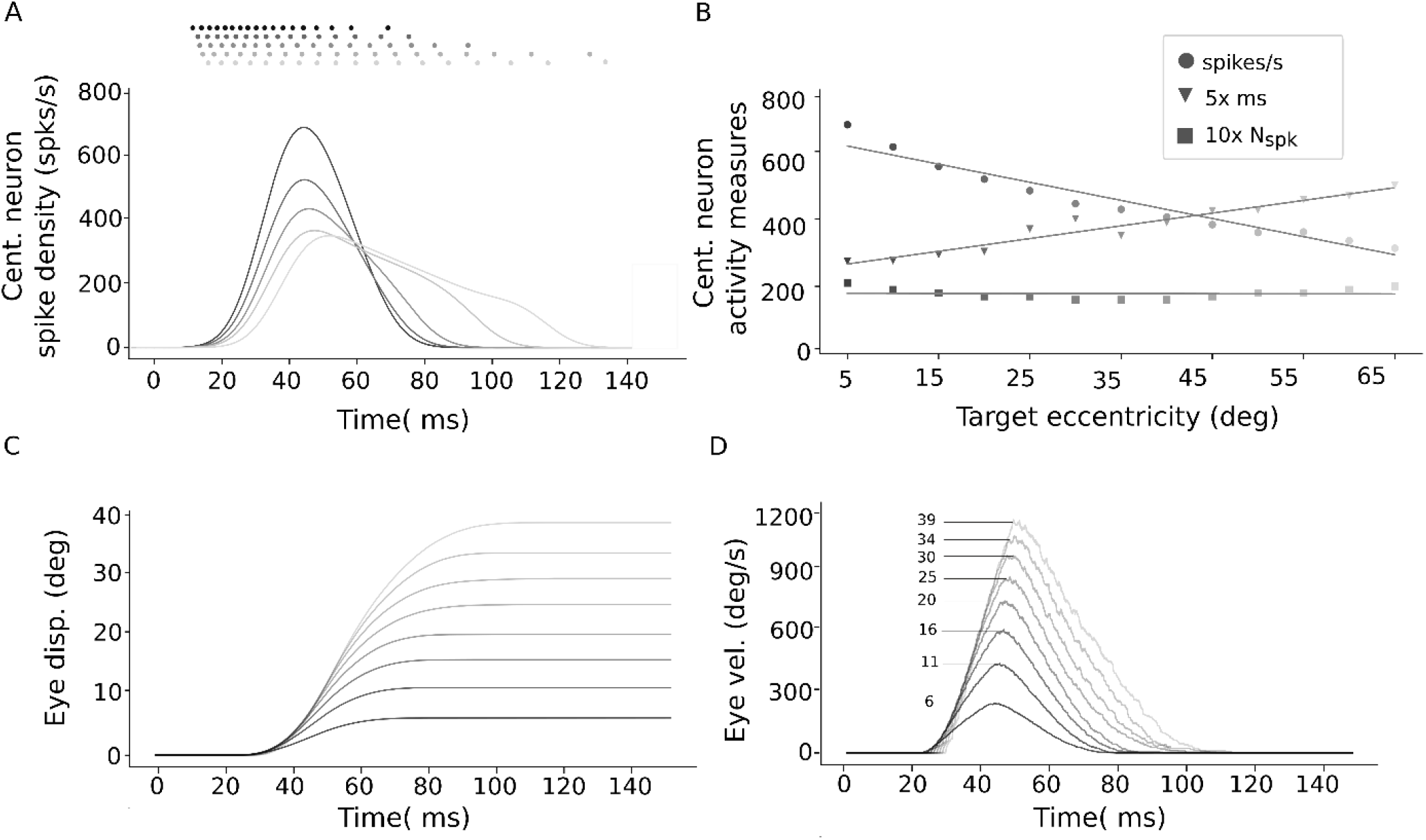
Results for default input. (A) Spike trains (top) and spike-density bursts profiles for five different central cells of the SC populations, in response to stimulation with the default input current at 5 different input sites (T=5,20,35,50,60 deg). Note the clear decrease of the peak firing rate, and increase in burst duration, with target eccentricity. The number of spikes in the burst varied slightly, but not systematically, between 16-19 spikes. (B) Central cell peak firing rates (dots) in spikes/s, burst duration (triangles), and the number of spikes (squares) for 13 default inputs applied between T = 5 and 65 deg. Note that the number of spikes for the central cell remains constant throughout the motor map, while the peak firing rate at the caudal sites drops to barely 50% of the rostral stimulation site. The burst durations of the central cell increase monotonically with the evoked movement amplitude. (C) Eye-displacement traces for different targets, at T = 5,10, 15, …, 40 deg, in the input layer. (D) Eye-velocity profiles for the corresponding eye-position traces in (C) (saccade amplitudes indicated). Note the clear increase in saccade duration, and the associated saturation of peak eye velocity as function of saccade amplitude.

Fig 6A shows how the evoked collicular bursts of the central cells in the recruited population systematically reduce their peak firing rates, and increase their duration, with increasing saccade amplitude. Panel 6B quantifies these properties as function of target eccentricity. According to the linear ensemble-coding model (Eq. 1), these burst properties underlie the nonlinear kinematic main-sequence properties of the associated saccadic eye movements. The saccades and their kinematics are illustrated in panels 6C, D, showing the evoked saccades for eight stimulation sites in the motor map. Note that the saccade duration increased with the saccade amplitude, and that the peak eye velocity increases in a nonlinear way with saccade size. A further characteristic property of the saccades concerns the skewness of their velocity profiles. Typically, the time-to-peak velocity of saccades changes little across a wide range of saccade amplitudes, while the saccade duration increases by saccade amplitude.

### 3.4 Effect of spatial-temporal changes in the input population

Fig 7 shows the collicular bursting profiles of the cells in the neural population for a saccade towards T=15 deg for input population sizes, *σ*_*pop*_, ranging from 0.05 – 1.0 mm, with *I*_*0*_ = 2.0 - 3.0 pA, and β = 0.03 s^-1^ (cf. Fig 2A). Fig 7A shows the peak firing rates of all recruited neurons in the SC layer for each stimulation condition (color encoded). The red curve corresponds to the default stimulation strength with *σ*_*pop*_=0.5 mm and *I*_*0*_ = 3.0 pA. The number of excited neurons, as well as their peak firing rates, increase with increasing input population size, saturating around the default stimulus condition at about 550 spikes/s for the central cell. In Fig 7B we show the normalized spatial-temporal activity patterns for the entire motor map for each of the different input populations. Note that the burst durations are the same for all stimulus conditions.

**Fig 7.**
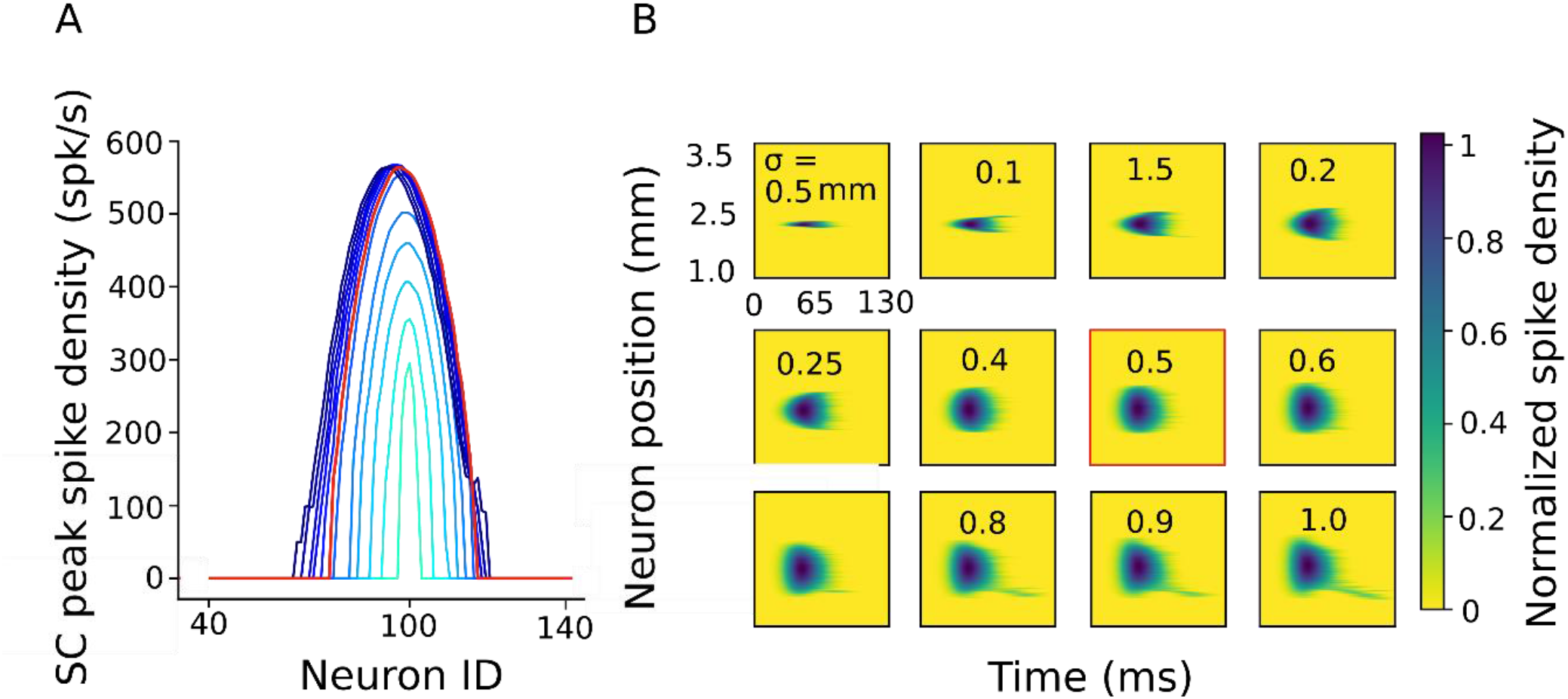
Burst profiles of the neurons in the network with lateral intra-collicular connections in response to the external input current that induces different input population sizes around T=15 deg. (A) Peak firing rates of the SC neural population. Red curve is the response to the default input current with σ_pop[_ = 0.5 mm. The population grows with input size up to the default current for low input strengths (light blue), after which it remains approximately invariant (dark blue). (B) Firing rate distributions of the neural population in the motor map as function of time, normalized to the absolute firing rate of the central cell (550 spks/s). Each panel shows the result for a specific input population size (indicated in mm). Red outlined panel corresponds to the default stimulation condition (0.5 mm). Panels preceding the default show that the SC population grows with the input population; panels following the default show that the number of active neurons remains approximately constant, even though the input population size grows to the double size.

Fig 8A, B shows the SC responses across the motor map while varying the duration parameter β of the input stimulus between 0.019 s^-1^ (long) and 0.03 s^-1^ (the default), for a fixed population size (*σ*_*pop*_ *=* 0.5 mm). The input current intensity, *I*_*0*_, co-varied with β between 1.2 – 3.0 pA in such a way that the number of spikes emitted by the input population decreased with increasing β (cf. Fig 2C, D, and the purple data points in Fig 2G, H). As a result, the total number of spikes emitted by the SC population, and hence the saccade amplitude, was independent of β (see also below).

**Fig 8.**
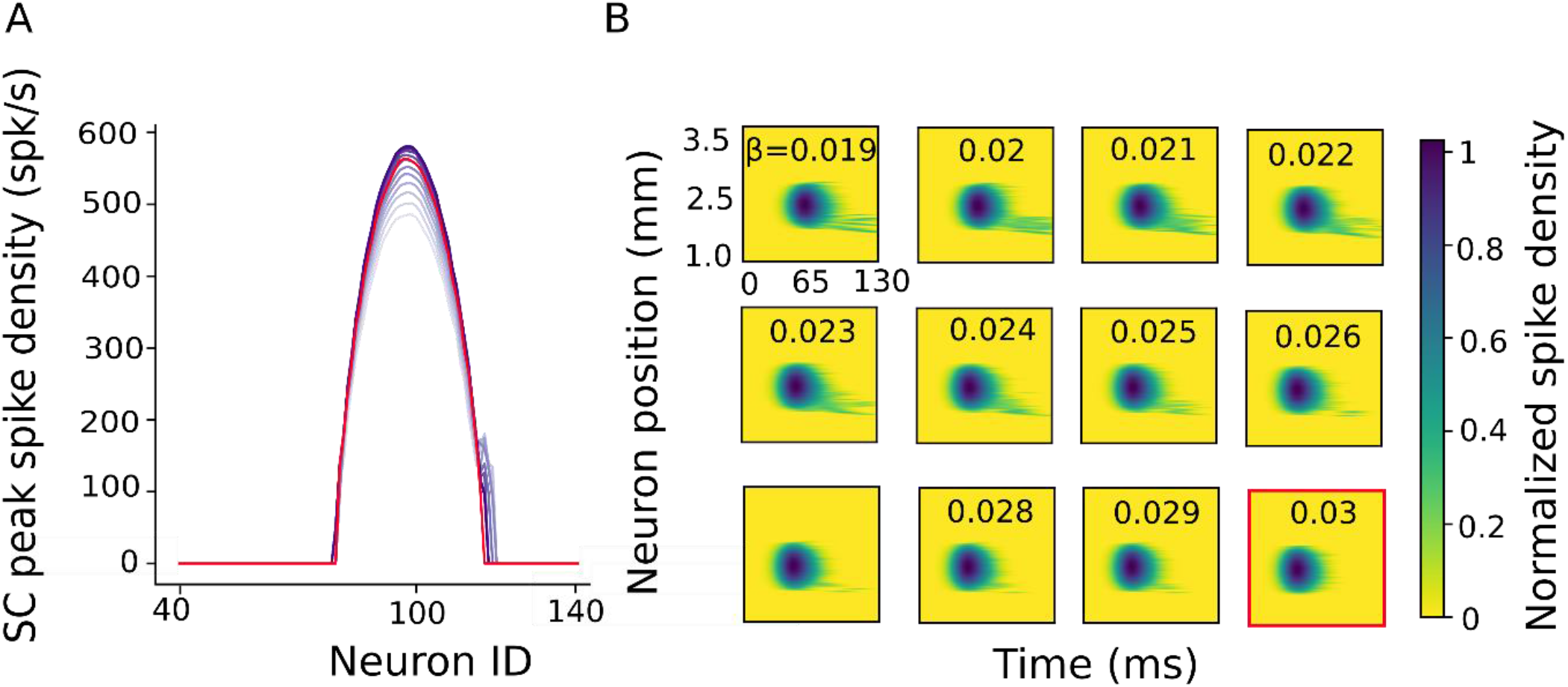
Burst profiles of the SC population in the network in response to the external input current with varying temporal properties (β, I_0_), selected such that the total number of input spikes sent to the SC motor map decreased with increasing β (see Fig 2C, D). Same format as in Fig 7. (A) Peak firing rates of the neural population in the motor map. Red curve corresponds to the default (β = 0.03 s^-1^). The SC peak firing rate increases with β and reaches a plateau around the default. The total number of SC spikes remained constant (see also Fig 2G, H, purple data). (B) Firing patterns of the neural population as function of time for the different β values. Note that burst durations decrease with increasing β.

### 3.5 SC output in response to spatial-temporal changes in the input

Fig 9 shows how the total number of spikes and recruited population size in the SC layer vary as function of the input stimulation parameters. Fig 9A shows the responses of the SC output layer when the size of the input population changes between 0.05 – 1.0 mm for target locations at T=10, 20, 30, 40 deg. For all four sites, the number of SC spikes and the SC population size show a saturating behavior: both output values are clearly lower than the default response (red dots) for small input population sizes and intensities. When the input exceeds the default values, the SC output saturates. Fig 9B, C shows the SC output behavior when the input current has a fixed population size of 0.5 mm but varies in its temporal behavior. In Fig 9B, β and I_0_ co-varied in such a way that the total SC output remained constant (see also Fig 2C, D). In Fig 9C, the variation in stimulation parameters kept the total number of spikes of the input layer constant (as in Fig 2E, F). Also, for these stimulation conditions the size of the SC population remained constant, but the total number of spikes from the population decreased with β for all four sites.

**Fig 9.**
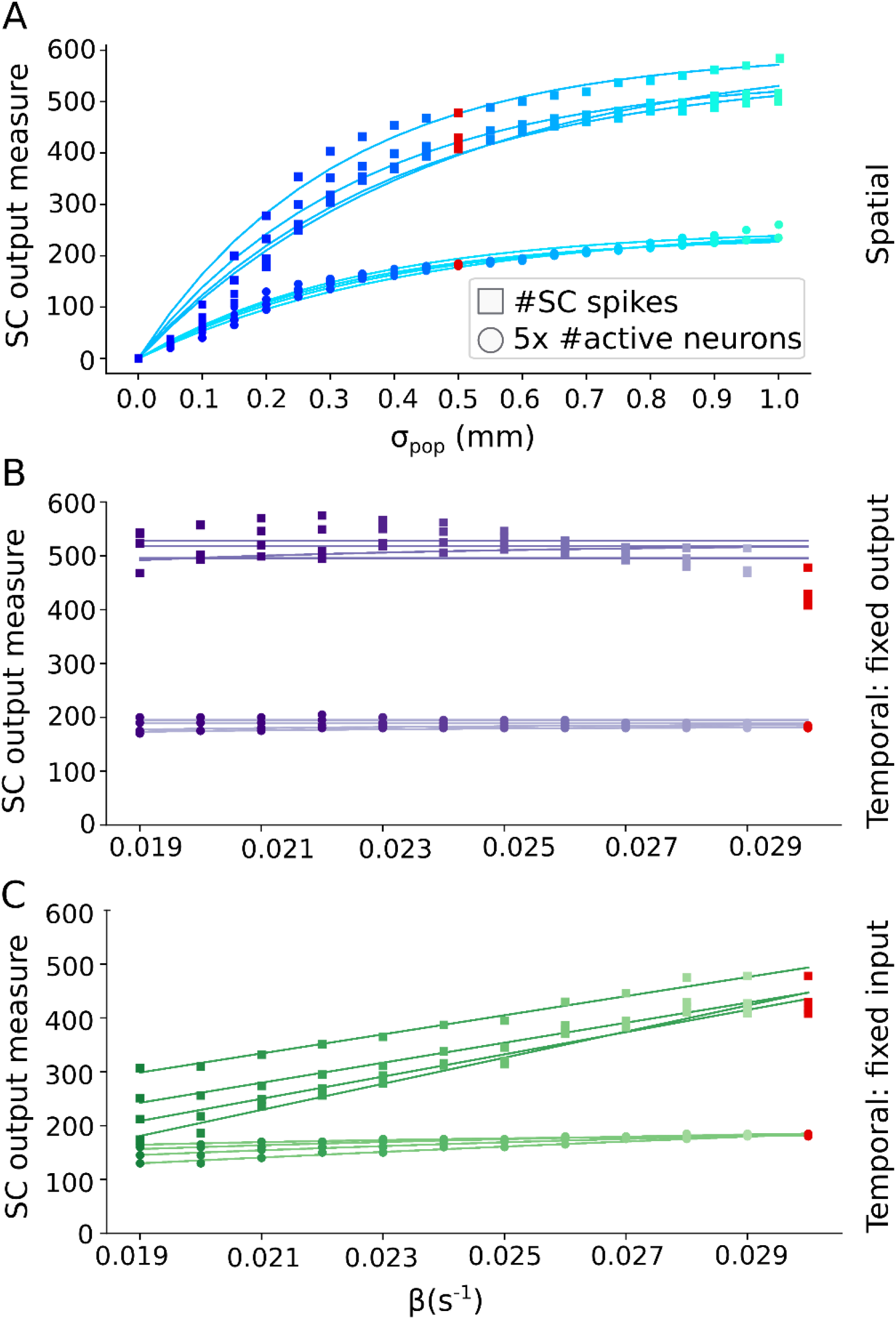
Burst properties of the SC saccade-related cells at 5 different sites (T=10,20,30,40 deg) as function of the input current parameters. (A) Number of active SC neurons (dots) and the corresponding total number of spikes (squares) from the SC map as function of input population size. Red dots correspond to the default stimulation. Curves increase with population size, up to the default, after which they saturate. (B, C) Number of active neurons (dots) and the corresponding total number of spikes (squares) from the SC layer as function of β. In (B), the stimulation parameters ensured a fixed number of spikes at the SC output (cf. purple curves in Fig 2G, H). In (C) the parameters ensured a fixed number of spikes from the input layer (cf. green curves in Fig 2G, H).

### 3.6 Saccade kinematics

In Fig 10 we show the evoked saccade amplitude and its peak velocity, as function of the external input current’s population size (Fig 10A, B), and as function of β (Fig 10C, D). The input stimulation was applied at three different sites on the input map, corresponding to T = 15, 20 and 30 deg, respectively. When the input population size fell below the default value of *σ*_*pop*_ = 0.5 mm, evoked eye movements were smaller than the intended target location. Around the default size of 0.5 mm (red symbols), the evoked saccade amplitudes approached the final, site-specific values (Fig 10A). Above the default population size, saccade amplitudes remained at the site-specific size (Fig 10A) over the full range of input strengths. The associated peak eye-velocity followed a similar input-dependent behavior for changes in the input population size (Fig 10B). Fig 10 C, D show the eye displacement amplitude and peak eye-velocity as function of β. The input current has a fixed population size (0.5 mm) and variable duration and strength to generate a fixed number of SC output spikes. The evoked saccade amplitudes remained close to the site-specific optimal values for all values of β. Note that the peak velocity increases slightly with β: short input bursts (i.e., high-frequency input stimulation) yielded higher velocities than longer inputs (low-frequency stimulation).

**Fig 10.**
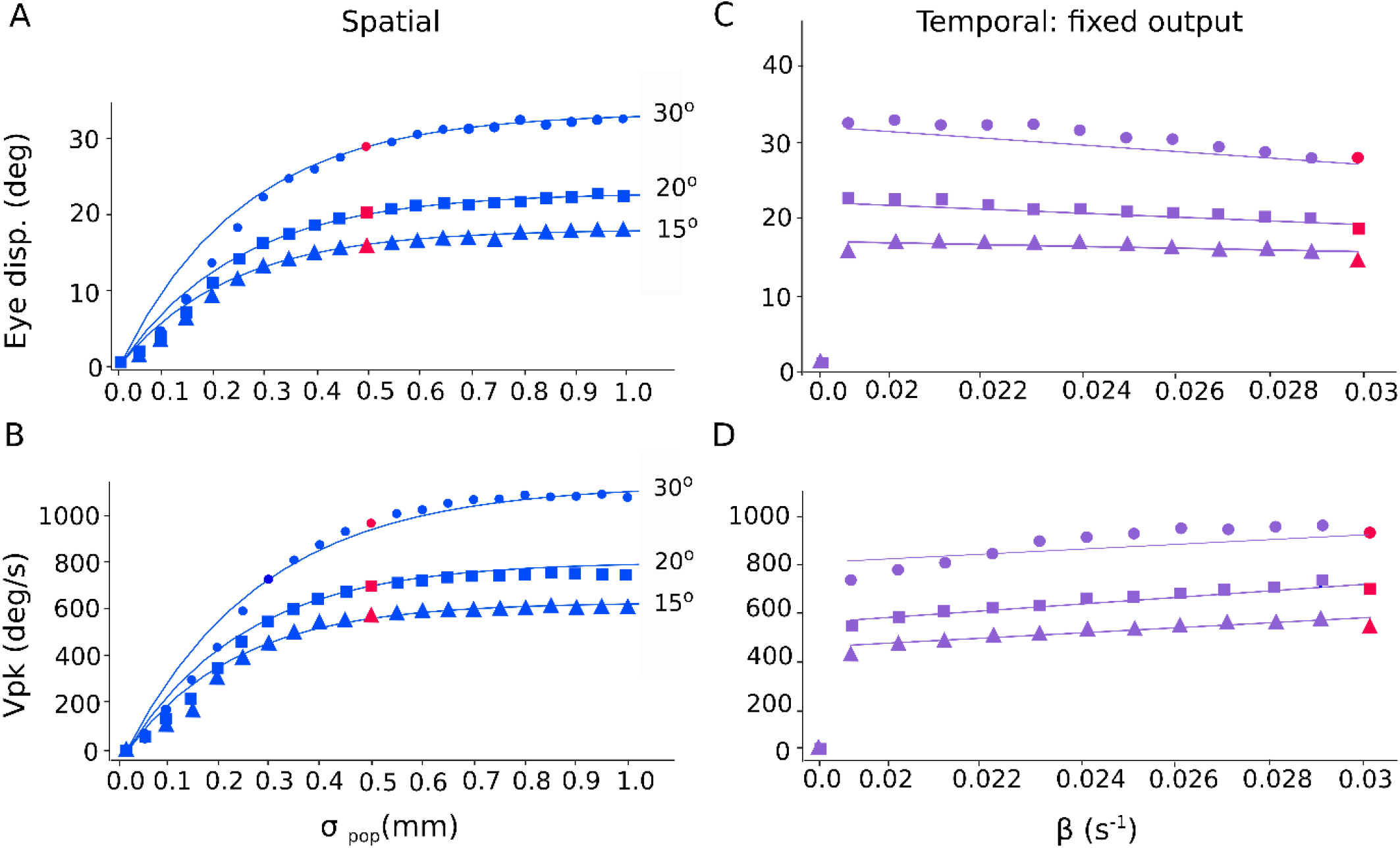
(A) Eye-displacement amplitude, and (B) peak eye velocity, as a function of the input current’s population size for stimulation at sites corresponding to T = 15, 20 and 30 deg. Beyond the default input population size of σ_pop_ = 0.5 mm (red symbols), the eye displacement amplitude and peak eye velocity are nearly independent of stimulation strength, while below 0.5 mm they both systematically decrease with decreasing input size. (C) Eye-displacement amplitude, and (B) peak eye velocity, as a function of the input current’s temporal parameter at sites corresponding to T = 15, 20 and 30 deg. The input currents generate fixed number of spikes at SC layer. Whereas the eye displacement remained invariant, peak eye velocity increased with β: the shorter the input duration (large β), the higher the velocity.

The kinematic main-sequence behaviors of the model’s saccades are quantified in Fig 11. The nonlinear amplitude-peak velocity relation of the model is quite comparable to the results from actual saccades, reported for monkey and human [2]. To quantify the model’s output in response to the default current stimulation applied at different input sites, we used a saturating exponential:

**Fig 11.**
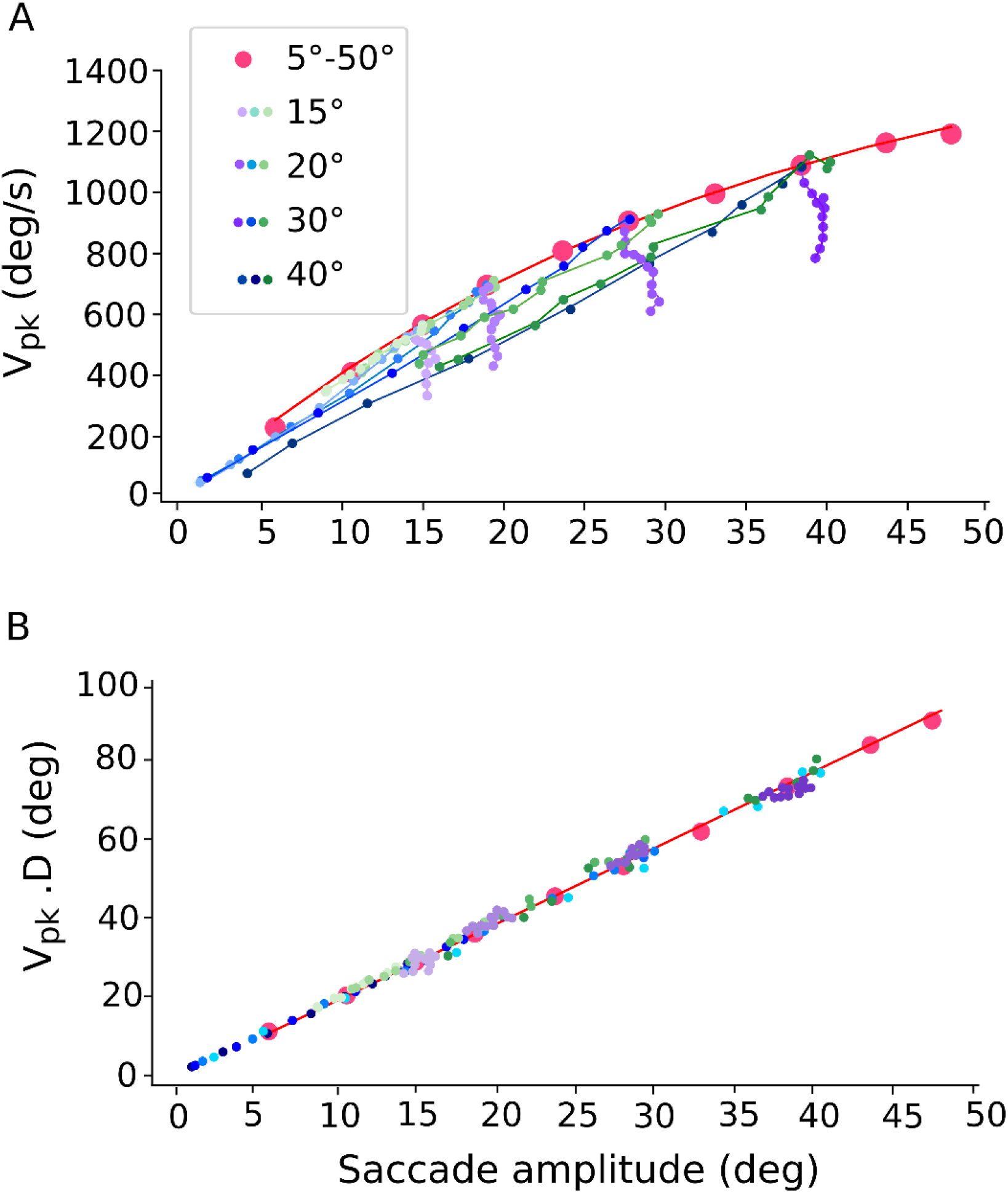
Nonlinear main-sequence behavior of the model. The red dots correspond to the output of the model for the default input strength applied at 10 input sites from 5-50 degs. (A) Red dots: Saturating amplitude-peak eye velocity relation (Eq. 8) for the default input current (σ_pop_ = 0.5 mm; β = 0.03; I_0_ = 3.0 pA) applied at 10 different sites. Blue dots: peak eye velocity of saccades evoked at sites T = 15, 20, 30 and 40 deg, respectively, for input currents with different input population sizes (from 0.05 – 1.0 mm), and fixed β. Purple dots: peak eye velocity of saccades evoked at the same sites, for input currents (σ_pop_ = 0.5 mm) with changing (β, I_0_), which kept the number of SC spikes constant. Green dots: same, for input currents with changing (β, I_0_), keeping the number of input-layer spikes constant. (B) The model saccades all follow the strict linear relationship (Eq. 9) for all stimulation conditions, and for all fast and slow saccades of panel (A).

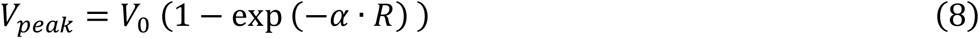

with V_0_ (deg/s) the saturation velocity at R=∞, and α (in deg^-1^) a measure for the slope of the relation. The red curve in Fig 11A corresponds to V_0_ = 1637 deg/s and α = 0.031 deg^-1^. Note that for the different input-stimulation conditions, the evoked saccade amplitudes could vary substantially (cf. Fig 10A), but the associated peak velocities of these smaller eye movements were also *slower* than for equally sized normal saccades, as all non-default data points fell below the default main-sequence curve. Thus, a fixed site in the SC motor map can generate saccades of different sizes, by variation of the recruited population size. In the model, the latter comes about by weak stimulation of the cells in the input layer.

The kinematics of pooled fast and slow saccades have been shown to be well described by the following linear relationship [2-3]:

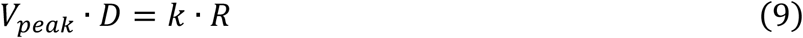

with *k* the slope (dimensionless). In Fig 11B we applied this relation to the default model saccades (red), obtaining a slope of *k* = 2.0, which is close to the experimentally obtained values for human saccades; it expresses the fact that saccades have single-peaked, skewed eye-velocity profiles that resemble a ‘triangular shape’ (cf. Fig 6D). Fig 11B shows that this relationship also describes all saccade data from the model, as also the smaller and slower eye movements evoked from the different input stimulation parameters all follow the same linear relationship.

## 4. Discussion

### Summary

We studied the properties of a simple, one-dimensional two-layer spiking neural network model with a cortical input and collicular output layer subjected to a large variation in the spiking input patterns. To investigate the relationship between the resulting SC firing patterns, saccade metrics, trajectories, and kinematics as function of the input current profiles, we varied the input stimulation patterns both in the spatial domain (input population size) and in the temporal domain (input population firing rates and burst durations).

Electrophysiological studies have shown that the saccadic system is quite robust against a large variability in SC input activation, but that at near-threshold stimulation levels the evoked saccades become both smaller and slower than expected from the normal main sequence [36, 38]. Furthermore, varying the input stimulation frequency, while keeping the total current fixed, modulates the saccade velocity [37]. Our previous spike-count model [40-41] could not account for these observations, as it was not designed to cope with a large spatial-temporal variation of the input.

By re-tuning the synaptic connectivity between the cortical input and SC output layers, and the intra-collicular excitatory-inhibitory lateral interactions, the new model was able to generate the changes in saccade properties that are associated with the variation in input population size and input firing frequencies, as obtained in electrophysiological studies.

### Mechanisms

Once SC neurons are recruited by the input, local excitatory synaptic transmission among nearby cells rapidly spreads the activation across the motor map to create a neural activity pattern, dictated by the most active central cells in the population. As a result, the burst shapes of the cells within the population were highly correlated [13,41]. Note that the evoked population activity in the SC output layer does not grow without bound, but is automatically constrained, both in its spatial extent, and in its bursting behavior (peak firing rates), by the inhibitory currents acting on the neurons whenever the external stimulation current attains high values. These inhibitory currents are due to the synaptic far-range lateral inhibition, which ensures that the population size remained within about 0.5 mm in diameter and was largely independent of the applied current when the stimulation parameters exceeded their default values. Conceptually, the lateral interactions normalize the population activity. In our updated network, the inhibitory and excitatory lateral connection strengths decrease (Fig. 3B) from the rostral to caudal zone by scaling parameter S_n_ thereby influencing the shape of the nonlinear relationship between saccade amplitude and peak eye velocity by reducing the firing rates of caudal neurons.

A systematic relationship between input current characteristics and the properties of evoked movements such as amplitude, velocity and duration has been demonstrated in electrical microstimulation experiments in monkey SC [36-38]. These studies reported that the evoked movement amplitude monotonically increased with the stimulation strength for low currents, while saturating at higher current strengths. These input-dependent properties had not been accounted for by the original linear ensemble-coding model [41], which assumed a fixed Gaussian input pattern, leading to a strong dependence on the input parameters. Yet, the actual electrophysiological results suggest that the external input acts predominantly as a trigger for the SC population-creating process. The intrinsic properties of the SC network subsequently set up the activity patterns of the cells, rather than the external stimulus itself. That is, the effects of electrical stimulation would be mainly caused by synaptic transmission, rather than by direct stimulation of the electric field.

A recently improved version of the SC model generated an SC population that relied less on the details of the input current, but the model could not produce small-amplitude, slow movements near the stimulation threshold. In addition, the input currents were described by stylized rectangular pulses, rather than by realistic spikes from cortical population inputs [40].

Here we improved the spiking-neural network model to generate small amplitude, slow saccades at low currents, increasing to a site-specific maximum at higher current strengths. We showed that the intra-collicular lateral connections could be tuned to generate saccades that faithfully followed the nonlinear main-sequence relations of normal, visually evoked saccades (Fig 8). Importantly, above the default value, the saccade metrics were unaffected by changes in the input stimulation parameters (Fig 10C). In addition, the saccadic peak eye velocity was modulated by the temporal properties of the input current: at short input burst durations (with a constant number of SC spikes), the evoked saccade velocities were higher, than at longer input burst durations (Fig 10D).

### Relation single-unit SC activity and ensuing saccade

Note that the linear ensemble-coding model predicts, in its simplest form, that the number of spikes of a given SC neuron for a fixed saccade should always be the same. However, this prediction hinges on the rather strong assumption that a single localized population of SC neurons generates the saccade, of which we can only be sure for a single-target visually evoked saccade in otherwise darkness and no other competing task demands or distractors. We applied this simple model also to a double-target stimulation task, which can yield a variety of double-step responses, including strongly curved saccades [45]. We showed that the expected activity for a given SC neuron could vary substantially, even when the overall vectorial displacement of the eye would be identical for all these trajectories. Thus, under such conditions, the strict relationship between SC spiking activity and saccade metrics and kinematics is broken, even though the saccade is still generated by linear ensemble coding.

In contrast, when these highly curved trajectories would result from an intended saccade to a single visual goal, but perturbed, e.g., by a blink response, the SC activity invariably relates to the overall saccade displacement vector, irrespective of the amount of curvature [46]. The reason for this discrepancy in neural behavior, despite an overall identical saccade behavior, is that when the total saccade trajectory results from the sum of two temporally overlapping sub-populations, related to the two goals, it can be generated in many ways. The problem with such situations is that with a single-unit recording technique it will be impossible to know the potential involvement of other parts of the SC motor map in a double-step scenario, beyond the cells around the recording electrode.

To avoid such ambiguities, we did not consider oculomotor scenario’s giving rise to several simultaneously activated cell populations, instead of a single localized (near-)Gaussian.

Peel and colleagues [35] observed that after local cooling of the FEF, the total number of spikes of an SC neuron for memory-guided saccades slightly decreased (by about 10% on average) when compared to visual evoked or pre-cooling memory-guided saccades, without affecting the overall saccade metrics. The decrease in spike count was absent for direct visual-evoked saccades. They proposed that the FEF-SC-Brainstem saccade pathway could be (acutely) bypassed by a parallel circuit (possibly involving the fastigial nucleus) to overcome the reduced input to the SC upon local FEF cooling. In this way, the extra signal from the parallel pathway would add to the reduced command from the SC, and still ensure a correct saccade amplitude. As in that case the relationship between the saccade metrics (number of spikes) and kinematics (firing rate) is broken, it may support the idea of a parallel pathway that can compensate for missing SC output. However, use of this alternative pathway is task dependent. Moreover, with the FEF intact, the strict spike-count – saccadic eye-displacement relationship holds for all saccades: slow memory-guided responses and fast direct visual-evoked saccades alike. Hence, under normal conditions, the direct FEF-SC-Brainstem pathway appears to be the major final common circuit for all saccades. This is also in line with the observation that an acute bilateral muscimol-induced inactivation of the SC practically abolishes the monkey’s ability to generate normal saccades [34].

### Future work

Although our improved model can account for a wide range of saccadic and SC response behaviors under widely different stimulation conditions, it still has several limitations. First, the model should be extended to two dimensions to enable saccades in all directions. The current model architecture allows for a relatively straightforward (but computationally expensive) extension and parameter tuning to a two-dimensional network [40, 47].

Recently, evidence was provided that in the head-unrestrained monkey the initial eye-in-head position strongly influences the gaze-shift kinematics, and that it has a systematic modulatory effect (‘gain field’) on the SC burst characteristics [48]. Interestingly, the large variation in gaze kinematics for a given gaze-displacement vector, was associated with a similar variation of the SC firing rates: slow gaze shifts were endowed with lower firing rates than fast gaze shifts. Yet, the instantaneous cumulative spike count of the SC cells faithfully encoded the straight gaze (i.e., eye in space) trajectory by following a similar linear relationship as was found for the head-restrained monkey’s eye movements [14].

A more complete model of the SC motor map in gaze control will have to include the control of eye-head gaze shifts, where the contribution of the eye- and head movement will systematically depend on the initial eye-in-head position, and in which an eye position signal itself is also able to modulate the spiking activity within the SC motor map.

Finally, an important factor that is still lacking in the current model is the presence of intrinsic multiplicative and additive noise in the parameters and neuronal dynamics, which would introduce additional variability in the evoked SC responses and the resulting saccades. These different aspects will be incorporated in our future studies.

## Acknowledgments

This work was supported by the European Union Horizon 2020 Programme, ERC Advanced Grant (2016) “Orient” 693400 (to A.A. and A.J.V.O.).

## Notes

### Competing Interest Statement

The authors have declared no competing interest.

## References

1. Bahill AT, Clark MR, Stark L. The main sequence, a tool for studying human eye movements. Math Biosci. 1975;24(3-4):191–204. https://doi.org/10.1016/0025-5564(75)90075-9

2. Van Opstal AJ, Van Gisbergen JA. Skewness of saccadic velocity profiles: a unifying parameter for normal and slow saccades. Vision research. 1987;27(5):731–745. https://doi.org/10.1016/0042-6989(87)90071-X

3. Evinger C, Kaneko CR & Fuchs AF. Oblique saccadic eye movements of the cat. Exp Brain Res. 1981; 41: 370–379. https://doi.org/10.1007/BF00238895

4. Van Gisbergen JA, Van Opstal AJ, Schoenmakers JJ. Experimental test of two models for the generation of oblique saccades. Exp Brain Res. 1985;57(2):321–336. https://doi.org/10.1007/BF00236538

5. Grossman GE, Robinson DA. Ambivalence in modelling oblique saccades. Biol Cybern. 1988;58(1):13–18. https://doi.org/10.1007/BF00363952

6. Smit AC, Van Opstal AJ, Van Gisbergen JA. Component stretching in fast and slow oblique saccades in the human. Exp. Brain Res. 1990;81(2):325–334. https://doi.org/10.1007/BF00228123

7. Van Gisbergen JAM, Robinson DA, Gielen S. A quantitative analysis of generation of saccadic eye movements by burst neurons. J Neurophysiol. 1981; 45(3):417–442. https://doi.org/10.1152/jn.1981.45.3.417

8. Scudder CA. A new local feedback model of the saccadic burst generator. J Neurophysiol. 1988; 59 (5):1455–1475. https://doi.org/10.1152/jn.1988.59.5.1455

9. Harris CM, Wolpert DM. Signal-dependent noise determines motor planning. Nature. 1998 ;394(6695):780–784. https://doi.org/10.1038/29528

10. Harris CM, Wolpert DM. The main sequence of saccades optimizes speed-accuracy trade-off. Biol Cybern. 2006;95(1):21–29. https://doi.org/10.1007/s00422-006-0064-x

11. Tanaka H, Krakauer JW, Qian N. An optimization principle for determining movement duration. J Neurophysiol. 2006;95(6):3875–3886. https://doi.org/10.1007/s00422-006-0064-x

12. Van Beers RJ. Saccadic eye movements minimize the consequences of motor noise. PloS one. 2008;3(4):e2070. https://doi.org/10.1371/journal.pone.0002070

13. Goossens HH, Van Opstal AJ. Optimal control of saccades by spatial-temporal activity patterns in the monkey superior colliculus. PLoS Comput Biol. 2012;8(5):e1002508. https://doi.org/10.1371/journal.pcbi.1002508

14. Goossens HH, Van Opstal AJ. Dynamic ensemble coding of saccades in the monkey superior colliculus. J Neurophysiol. 2006;95(4):2326–2341. https://doi.org/10.1152/jn.00889.2005

15. Van Opstal AJ, Goossens HHLM. Linear ensemble-coding in midbrain superior colliculus specifies the saccade kinematics. Biol Cybern.2008; 98(6):561–577 https://doi.org/10.1007/s00422-008-0219-z

16. Robinson DA. Eye movements evoked by collicular stimulation in the alert monkey. Vision Res. 1972;12(11):1795–1808. https://doi.org/10.1016/0042-6989(72)90070-3

17. Schiller PH, Stryker M. Single-unit recording and stimulation in superior colliculus of the alert rhesus monkey. J Neurophysiol. 1972;35(6):915–924. https://doi.org/10.1152/jn.1972.35.6.915

18. Sparks DL, Holland R, Guthrie BL. Size and distribution of movement fields in the monkey superior colliculus. Brain Res. 1976;113(1):21–34. https://doi.org/10.1016/0006-8993(76)90003-2

19. Ottes FP, Van Gisbergen JA, Eggermont JJ. Visuomotor fields of the superior colliculus: a quantitative model. Vision Res. 1986;26(6):857–873. https://doi.org/10.1016/0042-6989(86)90144-6

20. Moschovakis AK, Kitama T, Dalezios Y, Petit J, Brandi AM, Grantyn AA. An anatomical substrate for the spatiotemporal transformation. J Neurosci. 1998; 18(23):10219–10229. https://doi.org/10.1523/JNEUROSCI.18-23-10219.1998

21. Arai K, Das S, Keller EL, Aiyoshi E. A distributed model of the saccade system: simulations of temporally perturbed saccades using position and velocity feedback. Neural Networks. 1999;12(10):1359–1375. https://doi.org/10.1016/S0893-6080(99)00077-5

22. McIlwain JT. Lateral spread of neural excitation during microstimulation in intermediate gray layer of cat’s superior colliculus. J Neurophysiol. 1982;47(2), 167-178. https://doi.org/10.1152/jn.1982.47.2.167

23. Berthoz A, Grantyn A, Droulez J. Some collicular efferent neurons code saccadic eye velocity. Neurosci Lett. 1986;72(3):289–294. https://doi.org/10.1016/0304-3940(86)90528-8

24. Munoz DP, Pelisson D, Guitton D. Movement of neural activity on the superior colliculus motor map during gaze shifts. Science. 1991;251(4999):1358–1360. https://doi.org/10.1126/science.2003221

25. Waitzman DM, Ma TP, Optican LM, Wurtz RH. Superior colliculus neurons mediate the dynamic characteristics of saccades. J Neurophysiol. 1991;66(5):1716–1737. https://doi.org/10.1152/jn.1991.66.5.1716

26. Choi WY, Guitton D. Firing patterns in superior colliculus of head-unrestrained monkey during normal and perturbed gaze saccades reveal short-latency feedback and a sluggish rostral shift in activity. J Neurophysiol. 2009;29(22):7166–7180. https://doi.org/10.1523/JNEUROSCI.5038-08.2009

27. Jantz JJ, Watanabe M, Everling S, Munoz DP. Threshold mechanism for saccade initiation in frontal eye field and superior colliculus. J Neurophysiol. 2013;109(11):2767–2780. https://doi.org/10.1152/jn.00611.2012

28. Schiller PH, Tehovnik EJ. Neural mechanisms underlying target selection with saccadic eye movements. Prog Brain Res. 2005;149:157–171. https://doi.org/10.1016/S0079-6123(05)49012-3

29. Burman DD, Bruce CJ. Suppression of task-related saccades by electrical stimulation in the primate’s frontal eye field. J Neurophysiol. 1997;77(5):2252–2267. https://doi.org/10.1152/jn.1997.77.5.2252

30. Hanes DP, Patterson WF, Schall JD. Role of frontal eye fields in countermanding saccades: visual, movement, and fixation activity. J Neurophysiol. 1998;79(2):817–834. https://doi.org/10.1152/jn.1998.79.2.817

31. Sommer MA, Tehovnik EJ. Reversible inactivation of macaque frontal eye field. Exp. Brain Res. 1997;116(2):229–249. https://doi.org/10.1007/PL00005752

32. Dias EC, Segraves MA. Muscimol-induced inactivation of monkey frontal eye field: effects on visually and memory-guided saccades. J Neurophysiol. 1999;81(5):2191–2214. https://doi.org/10.1152/jn.1999.81.5.2191

33. Schiller PH, True SD, Conway JL. Effects of frontal eye field and superior colliculus ablations on eye movements. Science. 1979;206(4418):590–592. https://doi.org/10.1126/science.115091

34. Hepp K, Van Opstal AJ, Straumann D, Hess BJ, Henn V. Monkey superior colliculus represents rapid eye movements in a two-dimensional motor map. J Neurophysiol. 1993;69(3):965–979. https://doi.org/10.1152/jn.1993.69.3.965

35. Peel TR, Dash S, Lomber SG, Corneil BD. Frontal eye field inactivation alters the readout of superior colliculus activity for saccade generation in a task-dependent manner. J Comput Neurosci. 2020:1–21. https://doi.org/10.1007/s10827-020-00760-7

36. Van Opstal AJ, Van Gisbergen JA, Smit AC. Comparison of saccades evoked by visual stimulation and collicular electrical stimulation in the alert monkey. Exp Brain Res. 1990;79(2):299–312. https://doi.org/10.1007/BF00608239

37. Stanford TR, Freedman EG, Sparks DL. Site and parameters of microstimulation: evidence for independent effects on the properties of saccades evoked from the primate superior colliculus. J Neurophysiol. 1996;76(5):3360–3381. https://doi.org/10.1152/jn.1996.76.5.3360

38. Katnani HA, Gandhi NJ. The relative impact of microstimulation parameters on movement generation. J Neurophysiol. 2012;108(2):528–538. https://doi.org/10.1152/jn.00257.2012

39. Du Lac S, Knudsen EI. Neural maps of head movement vector and speed in the optic tectum of the barn owl. J Neurophysiol. 1990; 63: 131–146. https://doi.org/10.1152/jn.1990.63.1.131

40. Kasap B, Van Opstal AJ. Microstimulation in a spiking neural network model of the midbrain superior colliculus. PLoS Comput Biol. 2019;15(4):e1006522. https://doi.org/10.1371/journal.pcbi.1006522

41. Kasap B, Van Opstal AJ. A spiking neural network model of the midbrain superior colliculus that generates saccadic motor commands. Biol Cybern. 2017;111(3):249–268. https://doi.org/10.1007/s00422-017-0719-9

42. Goodman DF, Brette R. Brian: a simulator for spiking neural networks in python. Front Neuroinform. 2008;2:5. https://doi.org/10.3389/neuro.11.005.2008

43. Brette R, Gerstner W. Adaptive exponential integrate-and-fire model as an effective description of neuronal activity. J Neurophysiol. 2005;94(5):3637–3642. https://doi.org/10.1152/jn.00686.2005

44. Trappenberg TP, Dorris MC, Munoz DP, Klein RM. A model of saccade initiation based on the competitive integration of exogenous and endogenous signals in the superior colliculus. J Cogn Neurosci. 2001;13(2):256–271. https://doi.org/10.1162/089892901564306

45. Van der Willigen RF, Goossens HH, Van Opstal AJ. Linear visuomotor transformations in midbrain superior colliculus control saccadic eye-movements. J Integr Neurosci. 2011;10(03):277–301. https://doi.org/10.1142/S0219635211002750

46. Goossens HH, Van Opstal AJ. Blink-perturbed saccades in monkey. II. Superior colliculus activity. J Neurophysiol. 2000;83(6):3430–3452. https://doi.org/10.1152/jn.2000.83.6.3430

47. Kasap B, Van Opstal AJ. Dynamic parallelism for synaptic updating in GPU-accelerated spiking neural network simulations. Neurocomputing. 2018;302:55–65. https://doi.org/10.1016/j.neucom.2018.04.007

48. Van Opstal AJ, Kasap B. Maps and sensorimotor transformations for eye-head gaze shifts: Role of the midbrain superior colliculus. Prog Brain Res. 2019;249:19–33. https://doi.org/10.1016/bs.pbr.2019.01.006

